# A Novel Burn / Synovectomy Mouse Model for Temporomandibular Joint Osteoarthritis

**DOI:** 10.1101/2023.08.07.552340

**Authors:** Ginny Ching-Yun Hsu

## Abstract

Temporomandibular Disorders (TMDs) is present in 33% of the U.S. population. Currently available animal models do not faithfully simulate the native disease progression of TMJ OA. The initiation of TMJ OA requires both local trauma and systemic inflammation. In this study, we present a novel mouse model which reproduces these two conditions. This is achieved by a procedure involving both synovectomy (local trauma) and a distant burn injury (systemic inflammation). Its efficacy at inducing TMJ OA was assessed with histomorphology and radiographic evaluation at 1,3, and 9 weeks after the procedure. We found that burn/synovectomy mice demonstrated significantly more degenerative hard and soft tissue changes in TMJ than uninjured control mice or synovotomy mice. The observed histology in burn/synoectomy mice mimicked native TMJ OA disease progression in a faithful manner. This animal model is invaluable in future research of the mechanism and risk factors of TMJ OA.

## Introduction

Temporomandibular Disorders (TMDs) affect approximately 33% of the U.S. population^1^. Among these, Temporomandibular Joint (TMJ) Osteoarthritis (OA) accounts for pain symptoms in 10 to 17% of patients^2^. TMJ OA is characterized by synovitis, destruction of surrounding tissues and erosion of the subchondral bone with different level of pain and further result in facial asymmetry and jaw malfunction^3^. TMD is impacting the quality of life, and is associated with an annual healthcare cost of ∼$4 billion in the United States alone ^4^.

Animal models play a crucial role in understanding the complex TMJ OA disease progression and evaluating new therapeutic interventions. However, lack of a reliable *in vivo* model makes the research into the pathophysiology of TMJ OA is relatively restricted^5,6^. Besides genetically-induced animal models^7,8^, which could be time-consuming and expensive, and do not induce inflammation in the synovial capsule.

Intra-articular injection is a well-characterized preclinical model which induces OA by intra-articular inflammation, cytotoxicity, and direct cartilage damage ^5,6,9-14^. While it is easy to perform, its pathogenic mechanism differs from human TMJ OA and is only suitable for study of pain and treatment response ^15,16^. Other non-invasive means of inducing TMJ OA, such as mechanical loading, high-fat diet and sleep deprivation, only result in mild lesion and could be time-consuming. Could only mimic TMJ OA that arise from specific mechanisms ^11-13^.

In this study, we hypothesize that the concomitant use of two procedures, which include 1) synovectomy to cause local trauma, and 2) partial-thickness burn injury to cause systemic inflammation, can reliably induce TMJ OA in mice under an experimental setting. Through histologic examination and microCT imaging, we seek to validate our *in vivo* animal model in its ability to mimic natural TMJ OA disease progression and its value as a future research tool.

## Materials and Methods

This is a prospective controlled animal study approved by the Institutional Animal Care and Use Committee of Oregon Health & Science University (IP00004230). All animal procedures were carried out in strict accordance with good animal practices, as defined in the *Guide for the Use and Care of Laboratory Animals*.

A total of fifty-four 10-week-old mice were used for the current study, equally divided between three comparison groups: 1) burn/synovectomy group (n=18), 2) synovotomy group (n=18), and 3) control group (n=18). Within each group, there were three male mice and three female mice. The mice in group 1, the experimental group, underwent concomitant synovectomy and burn injury. The mice in group 2 received synovtomy alone. The mice in group 3 received no procedure.

### Surgical Technique

#### Anesthesia

The animals were anesthetized using 80 mg/kg of ketamine with 10 mg/kg of xylazine. 0.6 mg/kg of Buprenorphine Sustained-Release (Bup SR) were injected subcutaneously immediately prior to procedure.

#### Synovectomy / Burn

Before synovectomy, the left cheek was shaved with clippers and sterilized with povidone-iodine. A Y-shape incision will be made along the distal of the left eye to the left ear to expose the zygoma (Figure 1A), extending the incision approximately 0.5 cm to the ear canal so the TMJ can be easily visualized. The zygoma will be palpated underneath the temporalis and masseter muscles until the posterior end of the zygoma is reached and TMJ located distal to this, is identified (Figure 1B) and marked by the tip point of a #11 scalpel. Using the other hand to slightly move the mandible and cause condylar movement will further confirm the TMJ location. For synovectomy mice, after the incision the scalpel will be inserted 1-2 mm to the synovial capsule to increase the mobility of the TMJ (Figure 1C). The skin incision will be closed with a 5-0 vicryl stitches.

**Figure 1.**
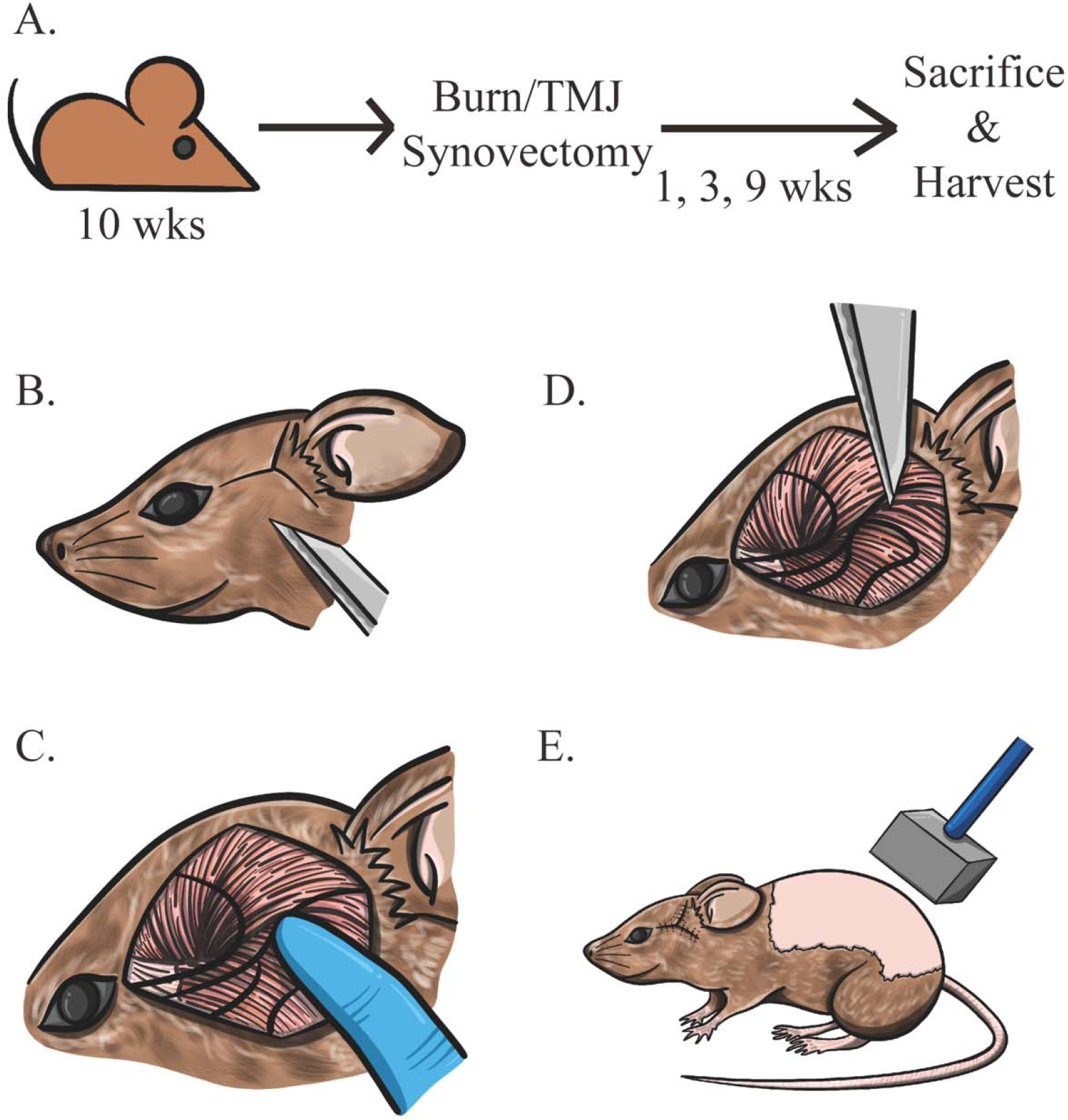
(A) Schematic illustration of Comparative Study Design: (B) Y-shaped incision to expose the zygoma extending from the lateral canthus towards the ipsilateral ear. (C) Identify the TMJ at the distal zygoma by palpation (D) At the posterior end of the zygoma, insert a #11 scalpel for 1-2 mm to make an incision in the synovial membrane and luxate the joint. (E) Perform the dorsal partial thickness burn.

During the same anesthesia session, a burn injury was made. The dorsal area from the spine to the left flank was shaved, exposing a skin area of at least 2 × 3 cm, and sterilized with povidone-iodine. Partial thickness burn injury was inflicted with an aluminum block weighing 35 g and measuring approximately 2 × 2 x 3 cm. The aluminum block was pre-heated to 60°C in a water bath, then applied to the shaved dorsum for 17 seconds (Figure 1D), using gravity to hold the block in place. This created an approximately 30% total body surface area burn in a 10-week-old mouse. Contact burn was chosen over flame or scald burn because it achieved a more uniform burn depth. The burn site was dried with gauze. Antibiotic dressing was applied.

#### Synovotomy

During synovotomy, a similar incision was made as described above for synovectomy, the tip of the #11 scalpel were made an incision on the synovial membrane. The skin was closed over the incision in a similar fashion.

#### Recovery

After the surgical procedure, both Group 1 and 2 mice received warmed resuscitation fluid with 0.5ml Lactated Ringer’s through intraperitoneal injection. The mice were given soft diet up to 3 days after the procedure. Subcutaneous injections of 0.6 mg/kg Buprenorphine SR were administered every 3 days.

#### Harvest

Within each group, six animals were sacrificed at 1, 3, and 9 weeks, respectively. They were euthanized with CO_2_ inhalation according to institutional guidelines. These time points were chosen to represent different stages in the inflammatory process. Deaths were ensured with cervical dislocation.

#### Outcomes

Our primary outcome is severity of intraarticular inflammation, as quantified by Osteoarthritis Research Society International (OARSI) score. Secondary outcomes include qualitative histologic exams and microCT imaging of cartilage changes. All specimens were analyzed in a blinded fashion.

After animal sacrifice, their TMJ were collected and fixed in 4% paraformaldehyde for 24□hours, decalcified with 14% ethylenediaminetetraacetic acid for 14 days, and embedded in OCT compound (Sakura) ^17,18^. Sagittal sections of the joint were sectioned to 6□μm thickness for histology. All images were obtained with Apotome 3 microscopy (Carl Zeiss Microscopy) ^19^ for qualitative histologic exam.

OARSI scoring were performed in a blinded fashion using medium coronal TMJ sections to represent the weight-bearing area. Safranin O/fast green stained sections were used to assess for cartilage injury. OARSI grading ranged from G1 to G6. OARSI score was calculated according to the formula: score□=□grade (G1 to G6)□×□stage (S1 to S4) ^20,21^. OARSI score ranged from 0–24. was determined according to the formula: score□=□grade (G1 to G6)□×□stage (S1 to S4) ^20,21^.

In order to quantify the subchondral bone volume changes after advanced OA, we performed microcomputed tomography (μCT) on specimens harvested at week 9. We defined the region of interest as the whole subchondral bone above the condyle neck. Using the Inveon high-resolution microcomputed tomography system (Siemens, Malvern, PA), images were acquired using a 1,440-step single projection pattern with 0.25 degrees arc separation. Projections were set at 80 kV and 500 μA with 610ms exposure/projection with a 0.5-mm aluminum filter. The charge-coupled device magnification was set to high and the field of view was 22.56 transaxial by 8.2mm axial. Binning was set to 0, resulting in a calculated 10.68 micron voxel size. Images were reconstructed with Inveon Acquisition Workplace software using the Feldkamp algorithm, no downscaling, slight noise reduction setting, Shepp-Logan filter, and using the mouse setting for beam hardening. Reconstructed images were converted to DICOM format and analyzed using ImageJ.

We defined the region of interest as the whole subchondral bone above condylar neck for 3-dimensional reconstruction and bone volume analysis.

### Statistical analyses

Ordinal variables like OARSI grade and scores were presented as medium (mininmum∼maximum). Statistical analyses were performed using two-way Mann-Whitney-U Test. Continuous variables like bone volume were presented as average +/- standard deviation. Two-way ANOVA followed by Tukey’s post hoc test. Values of p < 0.05 were defined as statistically significant. All statistical analyses were performed using GraphPad Prism 9.5 (GraphPad Software, Inc. San Diego, CA).

## Results

We were able to execute the above described procedure and specimen harvest with no deaths that occurred outside of planned euthanasia time points. Quantitative data from the three comparison groups are tabulated in Table 1.

**Table 1.**
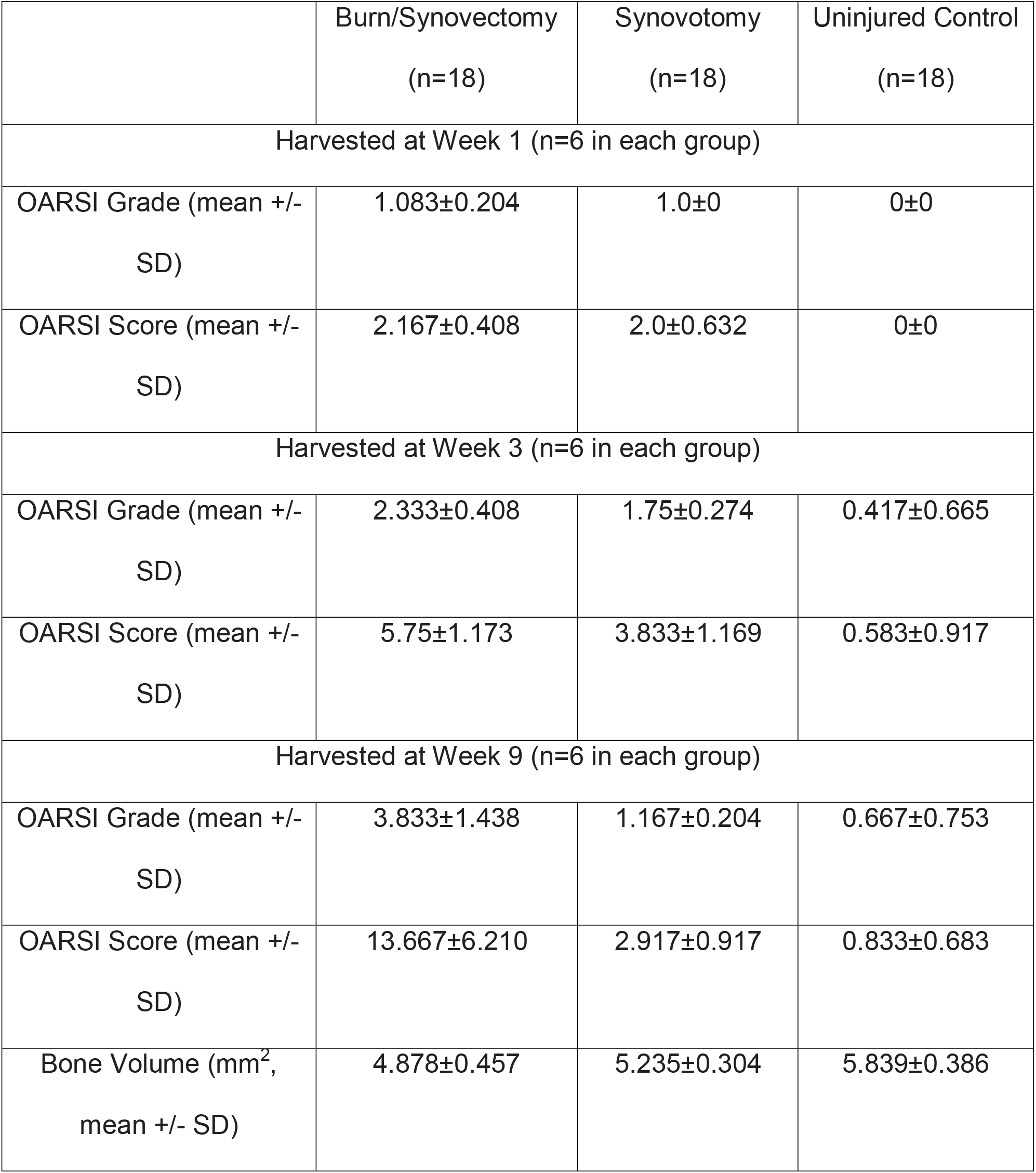
Summary of Radiographic and Histologic Findings on Various Groups

### Histologic Examination

We compared the severity of TMJ OA between burn / synovectomy mice and mice that received either just synovotomy or no injury by histologic examination (Fig 2). While the control mice remained normal, both burn/synovectomy and synovotomy mice displayed OA histologic features in their condyles as early as 1 week after surgery.

**Figure 2.**
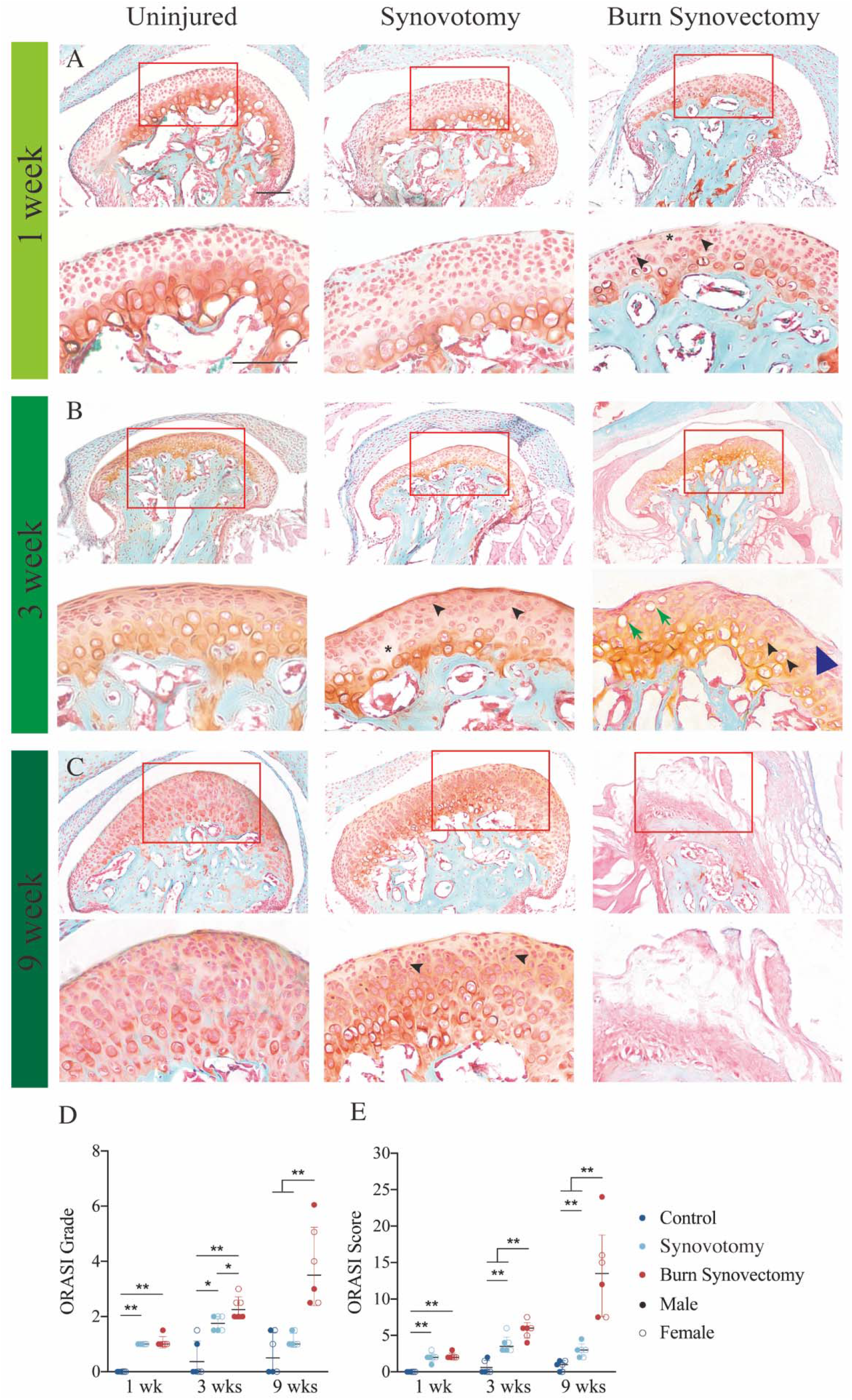
Histology Examination of TMJ Sections Histology of the TMJ sections stained with Fast Green and Safranin O Red and histopathology grade assessment. Uninjured (Left Column), synovotomy Middle Column) and synovectomy / burn (Right Column) mice were sacrificed after 1 (2A),3 (Middlw Row) (2B), 9 (Bottom Row) (2C) weeks for TMJ harvest. Within each row, low (upper panel) and high (lower panel)-magnification images allow detailed examination of the condyle morphology. Asterisk - acellular areas under the articular surfaces. Blue arrowhead: discontinuation of the condylar surface. Black arrowhead – loss of regular columnar structure among chondrocystes. Green arrow - apoptotic chondrocytes. Data are expressed as means, with dots representing data from individual animals. n = 6 (3 male and 3 female) animals per timepoint per condition. Osteoarthritis Research Society International (OARSI) grade (2D) scores (2E) were evaluated. Data were analyzed by statistical analyses were performed using two-way Mann-Whitney-U Test. *=P < 0.05, **=P < 0.01 indicate the significant differences within different timepoint. Groups with different letters indicate significant differences (P <0.05) between different timepoint. Scale bars = 200 (low magnification) and 50 (high magnification) μm.

These include multifocal decrease in cell count (asterisks) and the formation of chondrocyte clusters (black arrows) within the superficial zone(Fig 2A). At 3 weeks, burn/synovectomy mice displayed more severe OA than synovotomy and uninjured control mice, as evident by their more irregular superficial zone (blue arrowhead), increased hypertrophic and apoptic chondrocytes (green arrow), and increased chondrocytes clusters (black arrowhead) from superficial to zone of chondrocytes (Fig 2B). Synovotomy also showed increased of lesions compared to uninjured control mice. However, majority of the lesion are appeared at the superficial layer of the condyle. At 9 weeks, the degenerative changes continued to progress, resulting in calcified bone erosion and cartilage fissures (Fig 2C).

We used OARSI system to quantify these degenerative changes (Fig 2D, E), scoring the degree of inflammation based on cell morphology, cartilage integrity and subchondral bone involvement. At 1 week after surgery, the burn/synovectomy group and synovtomy group displayed similar levels of increased lesions compared to uninjured control (Fig 2D. E), signifying an intact surface with minimal clusters, involving less than 25% of the examined area. At week 3, the lesions in burn/synovectomy was significantly increased compared to synovotomy and uninjured control groups in grade and score. Synovotomy also appear significantly increase pathology lesions compared to uninjured control. Interestingly, burn synovectomy group showed significantly increased of lesion compared to synovotomy group in grade but not in score. Indicating the tissue reaction were increased in burn synovectomy group compared to synovotomy group but not the joint involvement. At week 9, condylar destruction became even more pronounced within the burn/synovectomy group, with full depth erosion of the fibrocartilage (Fig 2C). Synovotomy group has significantly increased score but not grade compared to the uninjured group, denoting minimal lesion was identified but increased the joint tissue involvement in synovotomy group.

### Morphological analyses of TMJ OA

2D Micro-computed tomography (μCT) images of specimens harvested at week 9 were obtained for all three comparison groups (Figure 3). Normal joint morphology was found in the control group, where intact and uniform condylar subchondral bone was found (Fig 3A). While the synovotomy group displayed only limited subchondral changes (Fig 3B), the burn/synovectomy mice displayed severe defects (red arrow) on the condylar head (red arrowhead) (Fig 3C). Similarly, on 3D image reconstruction, control mice had smoother bone morphology, while craters were visible in the burn/synovectomy group.

**Figure 3.**
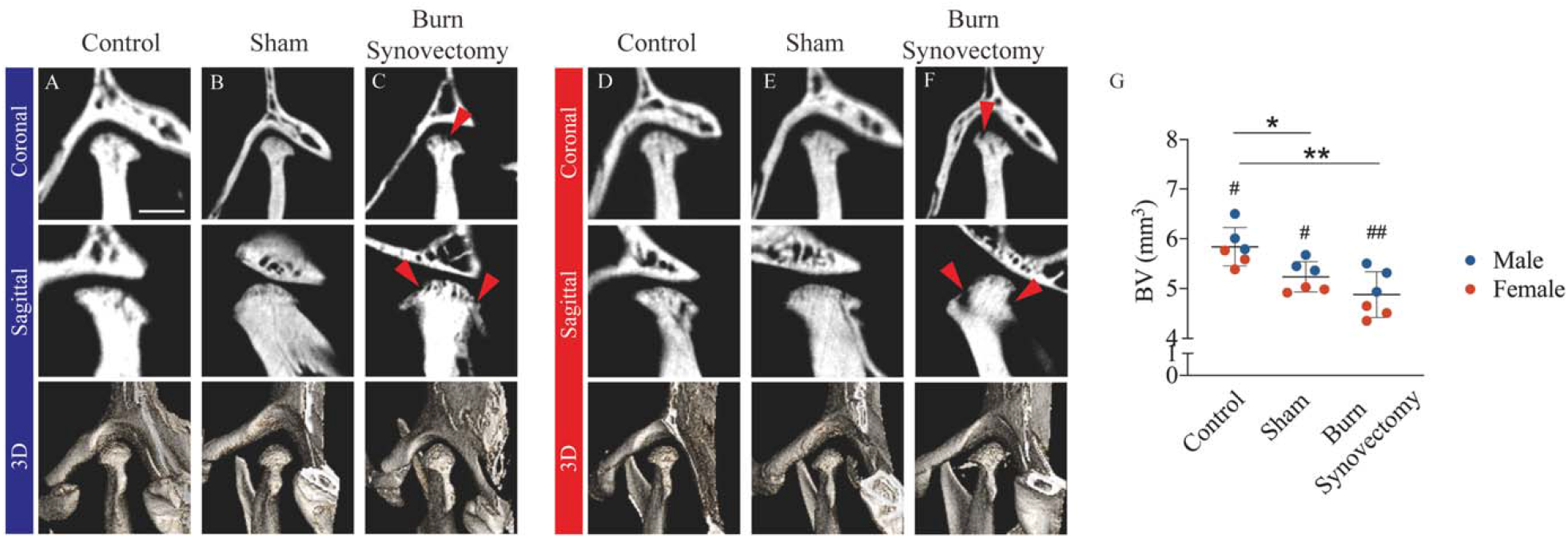
Radiographic changes in the condylar subchondral bone. Micro computed tomography(μCT) of the left TMJ of male (3A-C) and female (3D-F) animals after 9 weeks post-injuries. Representative Images of the coronal (upper), sagittal (middle) view and 3D reconstructions (lower) of left TMJs with uninjured control (3A,D), sham (synovotomy) (3B,E) and burn synovectomy (3C,F). Red arrow showed the bone erosion on the surface of the condyle. (3G) Micro-CT quantification of condyle bone volume. Data were analyzed by two-way ANOVA–repeated measures followed by Tukey’s post hoc test. *=P < 0.05, **=P < 0.01indicate the significant differences between different conditions and #=P < 0.05, ##=P < 0.01 indicate the significant differences between sex within same condition. Scale bars = 1mm

Severity of bone loss was quantified using bone volume analysis. Significantly higher bone volume was found in the synovotomy and control group than burn/synovectomy group, indicatng an increased level of erosion and subcondral bone turnover in the latter group. were discovered among burn/synovectomy group than either the synovotomy or the control group. Quantification analysis showed significant decrease in bone volume (indicative of erosion and turnover of subchondral bone) in the burn synovectomy group and sham group compared to uninjured control (Fig 3G). All three groups were evaluated between sexes. Male mice generally have increased bone volume compared to female counter part. However, the differences were increase in burn/synovectomy group.

## Discussion

In the current study, we developed an *in vivo* animal model that reliably simulates native TMJ OA disease progression. OA was induced by concomitant performance of a synovectomy, causing local traumatic injury, and a partial thickness burn, causing systemic inflammation. Our results showed that mice received burn/synovectomy exhibited more salient features of TMJ OA degeneration than mice that received either just synovotomy or uninjured mouse.

This is the first experimental animal model with a mechanism that imitates native TMJ OA. Since it is difficult to obtain TMJ samples from human patients, animal models play a key role for TMJ OA research ^5,16^. Our current study shows that a burn/synovectomy animal model well demonstrated the chain of cellular activities during TMJ OA progression. Start from the articular cartilage degeneration, manifested as over-production of proteoglycans and other extracellular matrix molecules, and the appearance of hyperplastic and proliferated chondrocyte clusters ^14^. Then, as the proteoglycans decrease on the surface of articular cartilage, the fibrocartilage layer fissures. Eventually, this degenerative sequence of events affects the subchondral calcified bone, leading to loss of bone volume, causing high bone turnover and aggressive osteoblast and osteoclast activities ^22,23^.

Another strength of our model is its reliability. We have observed similar severity and distribution of inflammation in between animals at each of the harvest time point. Our procedures allow easy standardization, as long as sterile technique was observed, and attention was paid towards uniform contact between the aluminum block and mouse skin. The primary limitation to our animal model lies in its translatability to human conditions. There is limited evidence in the literature to support the notion that systemic inflammation arising from distant burn sites increases the risk of TMJ OA. While TMJ OA is frequently observed in patients with craniofacial or cervical burn, it is unclear whether trunk or limb burn patients develop TMJ OA at a higher incidence ^24-26^. Furthermore, it is unclear if the systemic inflammation induced by burn injury is comparable to those from induced by other conditions, such as autoimmune diseases, with regard to its effect on the TMJ.

This is the first study that successfully simulates primary TMJ OA using a combined burn and synovectomy model, incorporating the effects of systemic inflammation to a disease process that was previously only simulated with mechanical injury. Due to its reproducibility and ease of operation, this model is ideal for research of the disease pathophysiology and treatment response.

## Disclosures

All authors state that they have no conflicts of interest.

## Acknowledgments

GCH is funded by NIH/NIDCR (K08DE031347) and American Association of Orthodontists Foundation.

## Notes

### Competing Interest Statement

The authors have declared no competing interest.

### Summary of Updates

The material/method and statistic has been revised. The result do not change the original finding but only optimized the quality of the manuscript

